# Leveraging unbiased LC-MS analysis to identify histone post-translational modifications associated with age-related decline of learning and memory

**DOI:** 10.64898/2025.12.05.692465

**Authors:** Shail U. Bhatt, Hasini Kalpage, Smaranda Bodea, Leticia Perez-Sisques, Ryan Gerkman, An Chi, M. Albert Basson, Sébastien Gillotin

## Abstract

Age-associated cognitive decline is a significant societal challenge, with several neurodegenerative diseases like Alzheimer’s disease becoming prevalent. Normal ageing is associated with several molecular, cellular, and metabolic hallmarks, including epigenetic mechanisms. In particular, dysregulation of post-translational modifications on histone tails is emerging as a mechanism associated with functional decline. The hippocampus is a region of the brain that is essential for the formation of episodic and spatial memories, which are particularly susceptible to age-related decline.

In this perspective manuscript, we have used mass spectrometry as an unbiased method to profile histone tail post-translational modifications both at baseline and after contextual fear conditioning (CFC) from young and aged mouse hippocampi to identify age- and activity-dependent changes.

Using this approach, we identify epigenetic marks not widely studied in ageing and in activity-induced learning. We propose a framework on how to integrate these epigenetic marks into current knowledge and discuss how to use this new analysis to formulate novel hypotheses that will broaden this field of ageing research.

We believe that this work will be of interest to scientists, clinicians and the wider public by making a significant contribution to our understanding of the molecular mechanisms responsible for age-associated cognitive decline and by reporting the identification of new mechanisms to focus on.

---

The cellular and molecular mechanisms responsible for cognitive decline with age remain incompletely understood. As a hallmark of ageing, epigenetic changes lead to irregular chromatin remodelling, aberrant DNA methylation, dysfunctional noncoding RNAs, and abnormal histone PTMs (1). Activity-induced PTMs have a role in synaptic plasticity and memory formation and are likely to work together with associated transcriptional processes for LTM formation (1, 2). Levels of histone acetylation, the most extensively studied PTM, decrease in ageing brains but is induced in rodent hippocampi by a learning stimulus like CFC (1). PTMs are diverse and mark a plethora of amino acid residues with many PTMs have not been studied in the context of fundamental and clinical research (2). Different PTMs can function antagonistically and synergistically. And these complex connections may be altered with age. However, the molecular associations between PTMs levels and age-related LTM formation have only been studied for a few specific PTMs. To link novel PTMs with hippocampal ageing and LTM-related mechanisms, we used an unbiased proteomic approach^1^ on histones extracted from 3- and 22-month-old mouse hippocampi (3), before and after being exposed to the CFC paradigm (Fig. 1A).

**Fig. 1.**
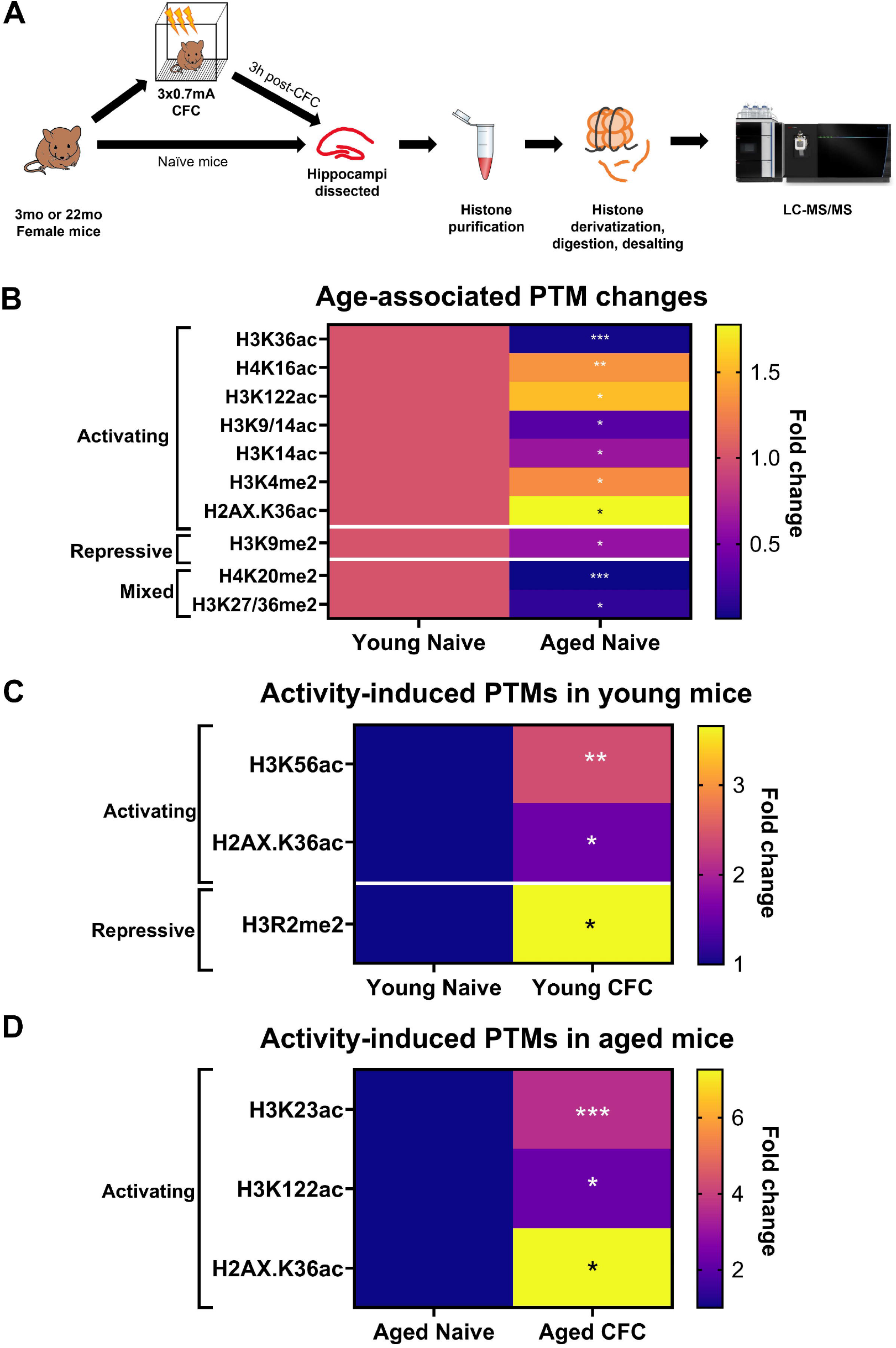
Significant histone PTMs altered with age and learning. A. Experimental design, 4 animals per condition. 3 hours post-CFC, all animals were sacrificed to dissect hippocampi and extract histones (Active Motif, cat #40025). Purified histones were derivatised with propionic anhydride, followed by trypsin/LysC digestion. Histone peptide N-termini were derivatised with propionic anhydride and desalted to be run through the Orbitrap Eclipse Tribrid Mass Spectrometer (MS). MS data was analyzed using Proteome Discoverer 2.5 with MS Amanda 2.0 and Sequest HT search engines. B. Heatmaps of PTMs modulated by age and segregated based on their effect on gene transcription (activating, repressive, or mixed/other effects), ranked by *P*-values from t-test (* p<0.05, ** p<0.01, *** p<0.001) and colour coded based on fold change over young naïve mice. C and D. Heatmaps of PTMs modulated by activity-dependent induction in (C) young and (D) aged mice with same analysis as in (B).

First, a comparison of PTM levels between the young and the aged naive groups identified 10 significant changes that included four acetylation marks (*i*.*e*. H3K9/K14ac, H3K14ac, H4K16ac and H3K122ac) (4, 5) all of which are known to be modulated in healthy ageing (Fig. 1B). We also identified several PTM changes for the first time, which included PTMs marking transcriptionally active genes, H3K36ac (decreased, *P-*value <0.0001), H2AX.K36ac (increased, *P-*value *=* 0.0391) and H3K4me2 (increased, *P-*value *=* 0.0327), the transcriptional repressive mark, H3K9me2 (*P-*value *=* 0.0437) and two PTMs with mixed functions, H3K27/36me2 (*P-* value = 0.0389) and H4K20me2 (*P-*value *=* 0.0005). Second, to identify activity-regulated PTMs, we compared hippocampal histones from naïve mice with histones from mice 3h after CFC (Fig. 1C, D). H2AX.K36ac was increased by CFC in young and aged mice (*P-*value = 0.0275 (Y), 0.0439 (A)), with a more pronounced increase in the aged CFC group (10-fold, Fig. 1D). Additionally, two marks were increased in the CFC young group:H3K56ac (*P-*value *=* 0.01) that marks transcriptionally active genes involved in newly replicated chromatin and H3R2me2 (*P-*value *=* 0.0282) that marks transcriptionally repressed genes (Fig. 1C). Likewise, two other marks associated with transcriptionally active genes were increased in the aged CFC group:H3K122ac (*P-*value *=* 0.0366) and H3K23ac (*P-*value *=* 0.000386) (Fig. 1D). Importantly, the modulation of each of these four PTMs following CFC was age specific.

Together this dataset constitutes a framework for exploring new mechanistic associations between these PTMs and molecular mechanisms underlying cognitive decline in ageing. Of all, H2AX.K36ac is the most compelling as it increased in all analyses (Figs. 1B-D). Like γH2AX, acetylation of K36 is known to be part of the IR-induced DNA damage response but unlike γH2AX its function to induce DNA repair is not fully explored (6). That is particularly relevant to study ageing and mechanisms associated with learning since respective accumulation of DNA damage and trigger of DNA breaks for gene transcription are well documented mechanisms for these two physiological processes (1). Therefore, testing whether H2AX.K36ac could be working in concert with the other identified PTMs also known to be linked with DNA damage and/or DNA breaks pathways would be a judicious hypotheses to test. For instance, could the simultaneous increases of H3R2me2 and H2AX.K36ac observed in our dataset be indicative of synergistic regulatory mechanisms to modulate different genes for CFC-induced memory formation in young animals? Likewise, could a concomitant decrease of H4K20me2 with an increase of H2AX.K36ac suggest dysregulation of DNA repair mechanisms in aged animals?

Histone PTMs are closely intertwined with chromatin compaction and chromatin topology which are both necessary for a dynamic gene transcription in learning processes. For instance, understanding if the decrease of H3K9me2 in aged hippocampi disrupts the normal organisation of heterochromatin could shed light on causal associations with impaired cognitive functions. Equally, increase of H3K56ac in young CFC mice but not in aged CFC mice could address whether nucleosome remodelling undergoes deterioration with age (7). Such hypotheses could be tested using LC/MS with purified nucleosomes alongside genome wide sequencing approaches.

Indeed, it will be crucial to identify where these PTMs are located in the genome of different cell types using chromatin immunoprecipitation-sequencing or Cut&Tag-sequencing after cell or nuclear sorting. Indeed, H3K9me2 and H3K14ac have been associated with synaptic plasticity and IEGs regulation, respectively (1, 2), but their contribution in the context of the ageing hippocampus is not fully established. Furthermore, it would be relevant to understand the elevation and subsequent activity-dependent induction of H3K122ac, known to be associated with oxidative stress and immune response that are key hallmarks of ageing. Finally, H3K23ac has been reported to play a role in learning and development (8) but its functional role in CFC-induced increase in aged hippocampi has never been explored.

By further deconvoluting the molecular mechanisms linked to the PTMs from this dataset, we can gain valuable insights into how these modifications influence the regulation of multiple genes in ageing. In return this would support the development of effective of strategies to prevent or reverse age-related decline. While we only identified methyl and acetyl groups among statistically significant PTMs, we detected other modifications (e.g., phosphorylation-Table 1) reinforcing the unbiased approach of using mass spectrometry. Several writers and erasers, including p300 and CBP, have been linked to neurodegenerative disorders, such as Alzheimer’s disease. Studying PTMs from this work in conjunction with selective knockdown of enzymatic proteins and pharmacological approaches in established models of AD (9) could be a rewarding approach to identify more selective therapeutics.

Ultimately, validating results in human cellular models and tissue while continuously integrating new PTMs into age predictor algorithms (10) will build a broader framework for dysregulated mechanisms in ageing and will ensure better translatability to the clinic.

## Methods

### Mouse models

Experiments were conducted using female C57BL/6J mice obtained from Charles River Laboratories. Young mice were 3 months old while aged mice were 18-22 months of age. Mice were group-housed under standard laboratory conditions with food and water ad libitum. All work involving mice was conducted in accordance with the UK Animals Scientific Procedures Act 1986 (PIL I44558708, and under the project licenses P8DC5B496//PP6246123 (Prof. Basson) and was approved by the King’s College London Animal Welfare and Ethical Review Board (AWERB).

### Contextual fear conditioning

Animals were placed in a soundproof fear-conditioning apparatus with stainless steel metal grid floor, containing a camera (Med Associates). To provide an olfactory cue, an ethanolsoaked tissue was placed under the grid in both training and testing sessions. For conditioning, mice were placed inside the chamber and left to freely explore it. After 148 s, three electric shocks (0.7 mA, 2 s each, 30 s apart) were administered. Mice were sacrificed 3 hours after the CFC.

### Histone extraction

Histone extraction was carried out using the column-based extraction protocol from Active Motif. Mice were sacrificed using cervical dislocation and the hippocampus was dissected. For nuclei isolation, fresh tissue was rinsed in ice-cold nuclei extraction buffer, was then minced into <1mm pieces and transferred to a homogenizer. Tissues were left overnight in the nuclei isolation buffer, in order to extract the crude histones. The crude histones were neutralized, and the samples were fed into a resin-packed column. Core histones were purified by running samples through the column and washing it with a series of proprietary wash buffers. Histones were then eluted, purified, and precipitated by adding perchloric acid at a final concentration of 4%. Samples were then washed thrice with 4% perchloric acid, acetone containing 0.2% HCl, and 100% acetone, respectively. Precipitated histones were instead dried using a speed vacuum.

### Mass spectrometry

Bottom-up MS/MS was used to elucidate post-translational modifications from the histone samples sent to MSD, as described by Sidoli and colleagues (2016). In brief, dry pellets were resuspended in 50mM NH_4_HCO_3_ (pH 8.0), and proteins were subsequently quantified. Histone derivatisation was carried out using propionic anhydride, followed by trypsin/LysC digestion of the samples. Histone peptide N-termini derivatisation was also carried out using propionic anhydride, after which all the samples were desalted, and run through the Orbitrap Eclipse Tribrid Mass Spectrometer, which made use of a 1-hour gradient (2-38% acetonitrile) as well as an EASY-Spray column. MS data was analyzed using Proteome Discoverer 2.5 with MS Amanda 2.0 and Sequest HT search engines.

## List of abbreviations

AD: Alzheimer’s disease
CBP: CREB-binding protein
CFC: contextual fear conditioning
DNA: Deoxyribonucleic acid
H2AX.K36ac: Histone 2A variant X lysine 36 acetylation
H3K4: histone 3 lysine 4 trimethylation
Me1: monomethylation
Me2: dimethylation
Me3: trimethylation
H3K9me: Histone 3 lysine 9 methylation
H3K14ac: Histone 3 lysine 14 acetylation
H3K27ac: Histone 3 lysine 27 acetylation
H3K27me: Histone 3 lysine 27 methylation
H3K36ac: Histone 3 lysine 36 acetylation
H3K56ac: Histone 3 lysine 56 acetylation
H3K122ac: Histone 3 lysine 122 acetylation
H3R2me: Histone 3 arginine 2 methylation
H4K16ac: Histone 4 lysine 16 acetylation
H4K20me: Histone 4 lysine 20 methylation
IR: Ionizing radiation
LC/MS: Liquid chromatography/mass spectrometry
LTM: Long-term memory
PTMs: post-translational modifications
RNA: Ribonucleic acid

## Footnotes

1 The protocol employed in this work involves chemical derivatization of histones with propionic anhydride, which affords improved coverage of multiple histone modifications. Due to the limited sample amounts we extracted from mice, it was not possible to perform an alternative derivatization with phenyl isocyanate, known to improve detection of lysine trimethylation modifications. We thus decided to prioritize the propionic anhydride derivatization reagent to afford better capture of novel or understudied histone marks.

